# Dual control exerted by dopamine in blood-progenitor cell cycle regulation in *Drosophila*

**DOI:** 10.1101/2021.03.29.437463

**Authors:** Ankita Kapoor, A. Padmavathi, Tina Mukherjee

**Affiliations:** Institute for Stem Cell Science and Regenerative Medicine (inStem), GKVK Post, Bellary Road, Bangalore 560065, India

**Keywords:** proliferation, cell cycle, hematopoietic progenitors, dopamine synthesis, dopamine receptor (*Dop2R*), dopamine transporter (*DAT*)

## Abstract

In *Drosophila*, definitive hematopoiesis occurs in a specialized organ termed “lymph gland”, where multi-potent stem-like blood progenitor cells reside and their homeostasis is central to growth of this organ. Recent findings have implicated a reliance on neurotransmitters in progenitor development and function however, our understanding of these molecules is still limited. Here, we extend our analysis and show that blood-progenitors are self-sufficient in synthesizing dopamine, a well-established neurotransmitter and have modules for its sensing through receptor and uptake via, transporter. Modulating their expression in progenitor cells affects lymph gland growth. Progenitor cell cycle analysis revealed an unexpected requirement for intracellular dopamine in the progression of early progenitors from S to G2 phase of the cell cycle, while activation of dopamine-receptor later in development regulated progression from G2 to entry into mitosis. The dual capacity in which dopamine operates, both intra-cellular and extra-cellular, controls lymph gland growth. These data highlight a novel and non-canonical use of dopamine as a proliferative cue by the myeloid-progenitor system and reveals a functional requirement for intracellular dopamine in cell-cycle progression.

## Introduction

Neurotransmitters are well appreciated for their roles in the central and peripheral nervous system to enable motor coordination, cognition and behavior control. However, an understanding of their roles outside this niche is gaining attention only recently. The immune system is one example that exhibits reliance on neurotransmitters for its development and function (Franco et al., 2007, Kenny and Ganta 2014, Hodo et al., 2020, Chen C et al., 2021). Neurotransmitter based regulation of hematopoiesis has been demonstrated in both invertebrates and vertebrates thereby suggesting a common axis employed by brain and blood system (Spiegel et al., 2008). Identification of neurotransmitters that are relevant for the development of blood cells and the mechanisms employed by these specialized molecules to coordinate hematopoiesis forms the central focus of our investigation.

In vertebrates, our understanding of neurotransmitters is from the stand-point of the bone marrow, where the hematopoietic stem and progenitor cells (HSPC), that generate differentiated blood cell types, are cradled by the bone marrow niche microenvironment. This niche is extensively innervated by the nervous system which upon release of neurotransmitters, like the catecholamine members, regulate HSPC mobilization through norepinephrine signalling (Katayama et al., 2006) or the migration and repopulation capacity of CD34^+^ progenitor cells via dopamine and epinephrine signalling (Spiegel et al., 2007). In invertebrates, such as *Drosophila*, similar implications of neurotransmitters governing hematopoiesis has also been established (Shim et.al., 2013, Madhwal et.al., 2020). The blood cell niche located within the larval hematopoietic organ termed “lymph gland” although lacks innervation, senses neurotransmitters that are derived systemically from the brain. Through neurosecretory routes, neurotransmitters are released from the brain into the hemolymph and then sensed directly by the hematopoietic niche. Our past and recent work (Shim et.al., 2013, Madhwal et.al., 2020) have implicated neurotransmitter function both as a signalling ligand and as a metabolite with non-overlapping roles in lymph gland growth, blood progenitor maintenance and differentiation. These studies not only validate conservation of processes underlying blood development, they also reveal intriguing insights that were previously uncharacterized. The use of neurotransmitters as metabolites by the blood system (Madhwal et.al., 2020) and the physiology driving their function (Katayama et al., 2006, Madhwal et.al., 2020) are some examples that warrants further understanding of other neurotransmitters in hematopoiesis. In this regard, the simplicity of *Drosophila* model system, along with the conservation of the nature of hematopoietic process and transcription factors guiding this process makes *Drosophila* a preferable system (Lebestky et al., 2000). In this study we address the importance of dopamine during blood development and identify its production within blood cells and its utilization as a growth promoting cue. Its role both as an intracellular metabolite and as a signalling ligand emerge as necessary modulators of cell cycle control in blood-progenitor cells.

Hematopoiesis in *Drosophila* is a biphasic event with an early primitive wave that takes place in the embryo and a later definitive wave which takes place in the larvae and originates in the lymph gland (Evans et al., 2003). The lymph gland is a specialized hematopoietic organ which is compartmentalised into an inner medullary zone (MZ) that is progenitor enriched and outer cortical zone (CZ) which harbours the differentiating hemocytes. A group of 30 cells cluster together to form a signalling niche and reside at the posterior end of the primary lobe, called the posterior signalling center (PSC) (Jung et al., 2005). Development and maintenance of MZ blood-progenitor cells rely on signalling derived from these zones (Sinenko et al., 2009, Mandal et al., 2007, Krzemien et al., 2007, Mondal et al., 2011) and also integrates systemic cues that are of nutritional and neuronal in origin. Of the systemic cues, the importance of neurotransmitter family of molecules have recently gained attention in progenitor development. GABA (gamma-aminobutyric acid) is one molecule, which is a well characterized neurotransmitter, but in the blood progenitor cells it has been shown to perform dual roles. GABA as a signalling entity activates GABA_B_R signalling and regulates maintenance of blood progenitor cells in an undifferentiated state (Shim et al., 2013). Our recent work highlights the use of GABA also as an intracellular metabolite whose catabolism stabilizes hypoxia inducible factor α (Hifα), protein, known an Sima in flies. Sima functions to maintain immune-competent progenitor cells necessary to respond to immune-challenges (Madhwal et al., 2020). While these findings have implied non-neuronal roles of neurotransmitters, they have highlighted the dependency of lymph gland progenitor cells on neurotransmitters. Specifically, the dual control exerted both as a signalling entity and as a metabolite, these small molecules emerge as integral regulators of lymph gland development.

Lymph gland development begins with the cessation of embryonic stage and with the onset of larval development, when at a first instar stage, the lymph gland comprises of single pair of lobes containing roughly 20 cells each. These cells proliferate to give rise to around 200 cells by the second instar stage (Jung et al., 2005). At this time, one also finds additional posterior lobes beyond the primary lobe. This growth of the primary lobe by 60 hours post egg-laying, is accompanied by the segregation of the MZ and CZ, which can be readily identified based on the markers expressed (Jung et al., 2005, Banerjee et al., 2019). The MZ progenitors express Domeless (Dome), a receptor of the JAK/STAT pathway, which serves as a progenitor marker and is detected until progenitor cells initiate differentiation following which they gradually lose Dome expression (Krzemien et al. 2010, Banerjee et al., 2019). The progenitors proliferate from early 1^st^ instar as evident by increased BrdU uptake and the presence of mitotically active cells (Jung et al., 2005, Krzemien et al., 2010). The progenitors eventually cease proliferation by third larval instar stage when they arrest in G2 phase of the cell cycle (Sharma et al., 2019) while the cells of the CZ initiate proliferation (Krzemien et al., 2010). The exit of progenitors from being in arrest into mitosis is linked to developmental signals perceived by them from differentiating cells (Mondal et al., 2011) which subsequently sets up the progenitors on a differentiation trajectory. Hence mechanisms governing cell cycle control in progenitor cells are also responsible for their maintenance and functionality. This journey of progenitors from the onset of proliferation at the late embryonic stage to reach quiescence by G2 arrest has intervening landmarks of cell cycle transitions that is not completely understood thus far.

In an attempt to address other neurotransmitters in progenitor development and function, a preliminary expression analysis of neurotransmitters and their sensing modules revealed components of dopamine pathway in progenitor cells. We explored dopamine in the development of myeloid-like progenitor cells during *Drosophila* larval hematopoiesis with the expectation to uncover any role dopamine might play in this process.

Our work demonstrates the utilisation of dopamine in regulating lymph gland size by controlling progenitor cell cycle phasing. We find that Dome positive progenitor cells synthesize dopamine and utilize distinct modules of dopamine synthesis, signalling and uptake to regulate lymph gland proliferative index. Early in lymph gland development, progenitor cells synthesize and internalize dopamine to coordinate exit of cells from the S-phase of the cell cycle, while signalling later in development through dopamine receptor function, *Dop2R*, drives progenitor cell exit from G2 into mitotic phase of the cycle. The dual regulation exerted by this monoamine contributes to maintenance of the cell-cycle phasing and building mitotic capacity of progenitor cells necessary for lymph gland growth. This regulation of size via dopamine is dispensable in the differentiating population i.e., the CZ thereby revealing specificity to dopamine’s requirement by the MZ progenitors. The study is the first description of dopamine in myeloid-progenitor proliferation and development. The importance of dopamine as a metabolite in induction of cellular proliferation portrays the non-canonical function of this monoamine in hematopoiesis.

## Results and Discussion

### The neurotransmitter dopamine is expressed in the growing lymph gland

The importance of neurotransmitters in *Drosophila* larval blood development has emerged from studies that have implicated GABA function in blood progenitor maintenance and differentiation (Shim et al., 2013, Madhwal et al., 2020) and serotonin in mature blood cells for phagocytosis (Qi et al., 2016). These findings demonstrate non-neuronal functions of neurotransmitters in mature differentiated cells as well as the development and maintenance of multi-potent blood progenitor cells. In this regard, dopamine is one monoamine whose role in mammalian immune function is known. Specifically, dopamine synthesis in the follicular helper T cells has been shown to enhance interactions with B cells to maximise the latter’s output (Papa et al., 2017). We undertook an antibody based staining approach to investigate if blood cells of the *Drosophila* larval lymph gland showed any levels of dopamine. We performed an analysis of this monoamine in a late 3^rd^ instar (96-102h <u>A</u>fter <u>E</u>gg <u>L</u>aying, AEL) lymph gland hematopoietic tissue, using an antibody raised against dopamine that is routinely used to detect dopamine in neuronal cultures (Banks et al., 2017, Chabrat et al., 2019) and also been reported in invertebrate hemocytes (Wu SF et al., 2015). The staining was undertaken in a genetic background that allowed comparative analysis of dopamine in progenitor cells (using a progenitor marker *domeMESO-Gal4, UAS-GFP*) which are Dome^+^ GFP^+^ as opposed to differentiating cells that are Dome^-^GFP^-^. As previously shown for GABA (Madhwal et al., 2020, See Materials and Methods for details), we used a stringent staining protocol (with 0.3% PBT (1X PBS with 0.3% Triton X 100) washes) to detect intra-cellular dopamine.

Interestingly, we detected dopamine in the 3^rd^ instar lymph gland (Fig. 1A-D, inset in B’-C’). To test the validity and specificity of the antibody in our assays, its localization was assessed in dopaminergic neurons in the brain that are positive for tyrosine hydroxylase (TH) expression. TH catalyses the first and the rate-limiting step in dopamine synthesis (Daubner et al., 2011). Neurons in the larval brain (Supplementary Fig. 1A-A’’, pointed in white arrows) that were positive for TH (Supplementary Fig. 1A’) overlapped with anti-dopamine antibody staining (Supplementary Fig. 1A’’). Apart from the intense anti-dopamine antibody expression in the TH^+^ neurons, the overall background staining with the antibody was negligible. This protocol validated the specificity of the dopamine antibody and the stringent washes ensured that the expression of dopamine detected in lymph gland blood cells was indeed intracellular.

**Fig 1:**
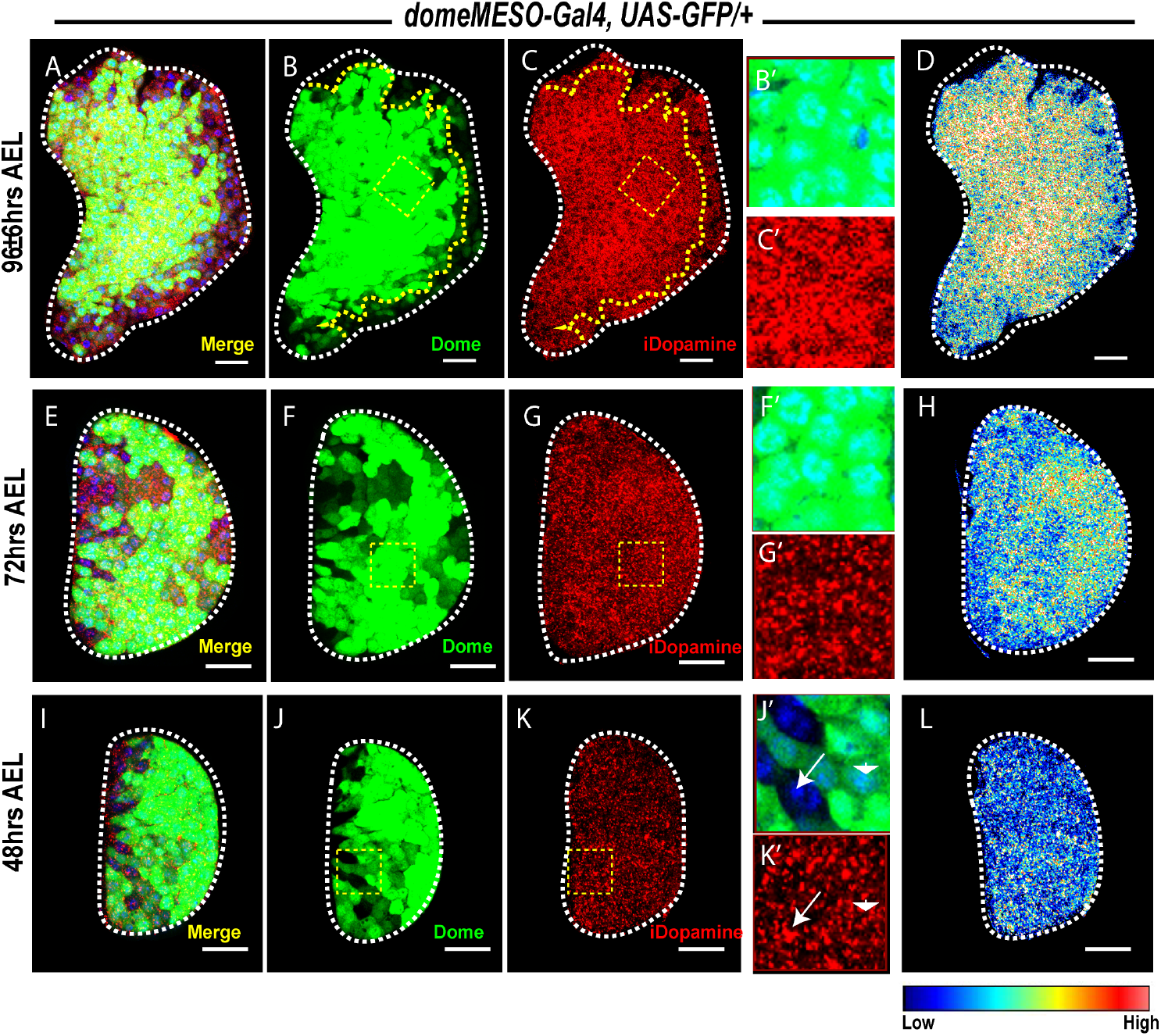
Dopamine expression during lymph gland growth. DNA is stained with Hoechst in blue, intracellular Dopamine (iDopamine) in red, *domeMESO* expression in green (representative of progenitors in the medullary zone, MZ). The iDopamine channel has been converted to spectral mode in panels D, H and L. All panels from A-C, E-G, I-K show a 20µm scale bar. The inset represents 2X zoom of region of interest (ROIs) marked in yellow dashed square. AEL indicates <u>A</u>fter <u>E</u>gg <u>L</u>aying. For better representation, the primary lobe of the lymph gland has been represented and outlined in white dashed lines. (A-D) Immunostaining with antibody against dopamine at mid-3^rd^ instar (96h AEL) shows an elevated expression of the molecule in the progenitors, marked with *domeMESO-GFP* and outlined with yellow dashed line as opposed to non GFP cells of the cortical zone. The inset shows Dome^+^ GFP^+^ cells (B’) and the corresponding dopamine levels (C’) in them. Similar expression analysis at early 3^rd^ (72h AEL, E-H) and early 2^nd^ instar (48h AEL, I-L) represents uniform dopamine expression pattern as evident from the inset in J’, K’ with arrowhead representing Dome^+^ GFP^+^ and arrow representing Dome^-^ GFP^-^ cells. The spectral mode of iDopamine representation in D and L shows elevated dopamine expression with increasing lymph growth.

We next conducted a temporal analysis of dopamine expression in lymph glands blood cells at earlier developmental time-point and observed that levels of dopamine detected at 72h AEL (Fig. 1E-H) was higher than levels at 48h AEL (Fig. 1I-L). Comparative analysis of dopamine levels detected at 48h AEL (Fig. 1K, L), 72h AEL (Fig. 1G, H) and 96h (96 Fig. 1C, D), implied increasing intracellular levels of dopamine over time. Moreover, as the animal progressed through the instars, dopamine levels were found to be significantly higher specifically in the Dome^+^ population later (Fig. 1C and G) in development than earlier (Fig. 1K). For better understanding of this observation, the dopamine channel was converted to spectral mode using ImageJ (LUT→NICE command). The blue pixels indicate lowest intensity, with the maximum depicted by red (See calibration bar below Fig. 1L). With this reference, it could be inferred that as the lymph gland increased in size, the dopamine levels concomitantly increased (Fig. 1L, H and D). We confirmed this differential expression of dopamine in the lymph gland zones by conducting the same staining protocol using cortical zone (CZ) marker (*Hml*^*Δ*^*-Gal4, UAS-2xEGFP*). A recapitulation of increased dopamine levels was observed when lymph glands from *Hml*^*Δ*^*-Gal4, UAS-2xEGFP* backgrounds were analysed at different developmental time points (Supplementary Fig. 1D, H, L and E, I, M). Hml or Hemolectin expression is detected is differentiating cells and in *Hml*^*Δ*^*-Gal4* background as well, Hml negative, undifferentiated progenitor cells showed high dopamine expression compared to Hml positive differentiating cells (Supplementary Fig. 1K’-L’). Thus further confirming the expression of dopamine in progenitor cells and ruling out the background effect of the genotypes utilised. Finally, dopamine levels were also detected in the Antp^+^ niche region and this showed no differences in intensity at the late 3^rd^ instar stage (110±6h AEL) compared to the remaining tissue (Supplementary Fig. 1N-P). Based on these observations we concluded that the neurotransmitter dopamine was detected in the lymph gland and gradually increased in progenitor cells as the tissue grew in size.

### Dopamine is synthesised by the medullary zone progenitors and regulates lymph gland size

Our observations on the detection of dopamine in the lymph gland progenitor cells led us to interrogate its source in this tissue. As the staining protocol utilised stringent detergent washes, the levels detected reflected residual dopamine present inside these cells. Therefore, the possibility of dopamine synthesis by the lymph gland cells was assessed.

Dopamine synthesis is a two-step process where the first step serves as the rate limiting step catalysed by the enzyme tyrosine hydroxylase (*TH*). The precursor molecule tyrosine is converted to an intermediate, L-dopa in this step which is then further converted to dopamine by the enzyme Dopa decarboxylase (*Ddc*) (Daubner et al., 2011) (Fig. 2A). We checked for the TH and Ddc enzymes in the developing 3^rd^ instar lymph gland (96h AEL) via immunostaining. We detected TH expression uniformly distributed in all cells of the lymph gland (Fig. 2B-D, inset in B’-C’). The expression of Ddc was elevated in progenitor cells (Fig. 2E-G). The presence of both the enzymes in the lymph gland blood cells implied a possibility of dopamine synthesis within the hematopoietic tissue. More importantly, elevated Ddc expression in progenitor cells along with elevated dopamine expression, implicated multi-potent progenitor cells as the site of dopamine synthesis (Fig. 2E-G, spectral mode in G and Fig. 1C-D). We therefore knocked down the enzyme’s *TH* and *Ddc* using a RNA*i* based approach in MZ progenitors using *domeMESO-Gal4, UAS-GFP*. This led to a substantial reduction in the levels of TH (Supplementary Fig. 2A-B’, quantification in E) and Ddc proteins respectively (Supplementary Fig. 2C-D’, quantification in F). The genetic knock-downs also led to a 50% reduction in intra-cellular dopamine levels in lymph gland blood cells when compared to levels detected in age-matched genetic controls (Fig. 2H-J, quantification in K). This analysis was undertaken at the wandering 3^rd^ instar larval stage (∼120h AEL), when the lymph gland has completely grown in size. In addition to the reduction in dopamine levels, a significant reduction in lymph gland size was also noticeable upon knocking down *TH* and *Ddc* expression in progenitor cells (Fig. 2L-N, quantification in O). This implied a functional role for intracellular dopamine synthesis in blood progenitor cells during lymph gland growth. We substantiated the small lymph phenotype by using another MZ specific *Gal4*-driver, *Tep4-Gal4, UAS-mcherry* to express *TH* and *Ddc* RNA*i* in progenitor cells (Supplementary Fig. 2G-I). *Tep4-Gal4*, driver, unlike *domeMESO-Gal4*, is expressed in a restricted subset of blood progenitor cells (Dey et al., 2016), but expressing the respective RNA*i*’s using this driver also led to a comparable reduction in lymph gland size (Supplementary Fig. 2J). This further confirmed dopamine’s role in lymph gland progenitor cells and its requirement in overall growth of the lymph gland.

**Figure 2:**
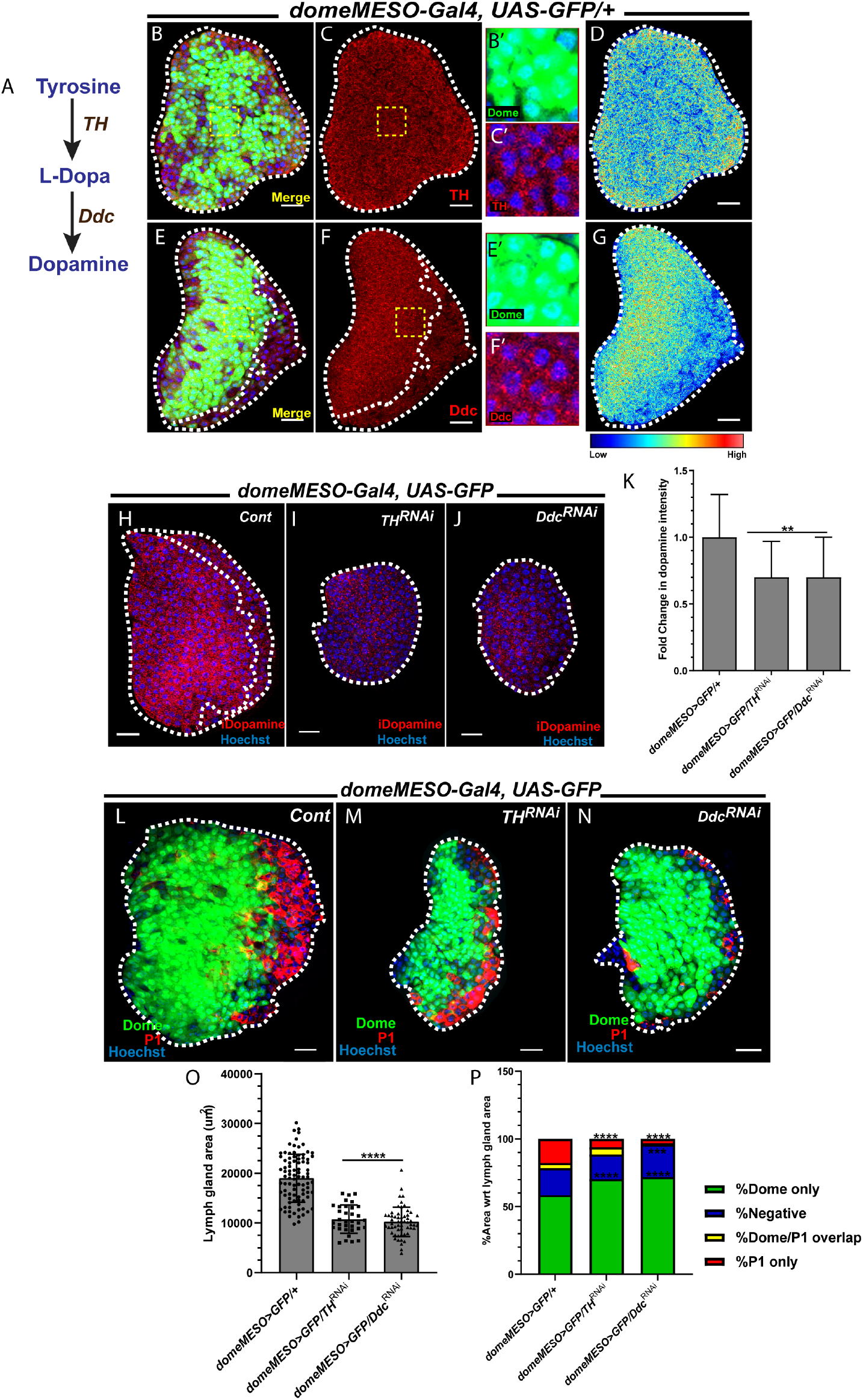
The medullary zone progenitors synthesize dopamine to control lymph gland size and differentiation. DNA is stained with Hoechst in blue, iDopamine in red, Tyrosine hydroxylase (TH) in red, Dopa decarboxylase (Ddc) in red, *domeMESO* (representative of progenitors) expression in green, P1 (representative of mature plasmatocytes) in red. All image panels show a 20µm scale bar. AEL indicates <u>A</u>fter <u>E</u>gg <u>L</u>aying. The quantifications in K and O represents mean with standard deviation and in P represents the mean. The statistical analysis applied in K is Mann-Whitney test while that applied in O and P unpaired t-test, two-tailed. ‘n’ represents the number of lymph gland lobes analyzed. For better representation, the primary lobe of the lymph gland has been represented and outlined in white dashed lines. (A) Schematic representation of the dopamine synthesis pathway where the amino acid tyrosine is converted to L-dopa via Tyrosine hydroxylase (*TH*), the rate limiting enzyme. Further L-dopa is converted to dopamine by the action of Dopa decarboxylase (*Ddc*). (B-D) The expression of TH in the mid-3rd instar (96h AEL) lymph glands of control (*domeMESO-Gal4, UAS-GFP/+*) shows a uniform cytosolic pattern (inset in C’) while the expression of Ddc in the similar stage and background shows elevated expression in the Dome^+^ GFP^+^ region (E-G), supporting the increased expression of dopamine in the progenitors. (H-J) Assessment of dopamine levels upon the knockdown of *TH* and *Ddc* in the progenitors using *domeMESO-Gal4, UAS-GFP* driver shows a 40-50% reduction in dopamine levels. (K) The fold change in dopamine levels in *TH* and *Ddc* loss of function in the progenitors. Compared to control (*domeMESO-Gal4, UAS-GFP*/+, n=40), a 0.5fold reduction in dopamine levels is observed in loss of *TH* (*domeMESO-Gal4, UAS-GFP*; *TH*^*RNAi*^, n= 17, **p=0.0017) and *Ddc* (*domeMESO-Gal4, UAS-GFP*; *Ddc*^*RNAi*^, n= 34, ***p=0.0003) (L-N) Loss of progenitor *TH* and *Ddc* function using *domeMESO-Gal4, UAS-GFP* shows a reduced lymph gland size compared to control (at wandering 3^rd^ instar, 120h AEL). (O) Quantifications of lymph gland size shows an almost 50% reduction in lymph gland size upon loss of *TH* (*domeMESO-Gal4, UAS-GFP*; *TH*^*RNAi*^, n= 34, ****p<0.0001) and *Ddc* (*domeMESO-Gal4, UAS-GFP*; *Ddc*^*RNAi*^, n= 56, ****p<0.0001) when compared to stage matched control (*domeMESO-Gal4, UAS-GFP*/+, n=91). (P) Quantifications of the proportions of % Dome (green bars for MZ), % negative (blue bar), %overlap (Dome and P1, yellow bars) and %P1 (red bars for CZ). An 8-10% expansion of Dome positive area was observed in the progenitor specific loss of *TH* (*domeMESO-Gal4, UAS-GFP*; *TH*^*RNAi*^, n= 15, ****p<0.0001) and *Ddc* (*domeMESO-Gal4, UAS-GFP*; *Ddc*^*RNAi*^, n= 18, ****p<0.0001) function w.r.t to control (*domeMESO-Gal4, UAS-GFP*/+, n=27). No significant changes in the % negative and % overlap areas were found upon *TH* knockdown (*domeMESO-Gal4, UAS-GFP*; *TH*^*RNAi*^, n= 15, p=0.4346 and p=0.0998, for %negative and %overlap respectively) compared to controls (*domeMESO-Gal4, UAS-GFP*/+, n=27). A mild reduction in % negative area was found upon loss of *Ddc* (*domeMESO-Gal4, UAS-GFP*; *Ddc*^*RNAi*^, n= 18, *p=0.0410) and a significant reduction in overlap of Dome/P1 population was noticed (*domeMESO-Gal4, UAS-GFP*; *Ddc*^*RNAi*^, n= 18, ***p=0.0003). The loss of dopamine synthesis in the progenitors caused a significant reduction in the plasmatocytes differentiation when compared to controls (*domeMESO-Gal4, UAS-GFP*/+, n=27, *domeMESO-Gal4, UAS-GFP*; *TH*^*RNAi*^, n= 15, ****p<0.0001 and *domeMESO-Gal4, UAS-GFP*; *Ddc*^*RNAi*^, n= 18, ****p<0.0001).

We assessed if progenitor-specific loss of dopamine synthesis (via *TH*^*RNAi*^ and *Ddc*^*RNAi*^) affected any aspect of progenitor maintenance or differentiation. This was addressed by assessing proportions of MZ and CZ in *TH* and *Ddc* loss-of-function backgrounds using respective markers of these zones (as described previously in Madhwal et.al., 2020). The assessment showed an expansion of around 8-10% in the Dome positive population in conditions with loss of both *TH* and *Ddc* from progenitor cells (Fig. 2P, green bars). This progenitor increase was associated with a concomitant reduction in the population of differentiated plasmatocytes that are positively marked for P1 (Fig. 2P, red bars). Compared to *TH, Ddc* loss of function showed stronger reduction in P1 differentiation (Fig. 2P, red bars). We checked for the crystal cell population through PPO staining (Evans et al., 2003) and observed that both *TH* and *Ddc* loss of function led to a significant reduction in crystal cell population (Supplementary Fig. 2L-N, quantification in O) as well. Similar analysis was undertaken using *Tep4-Gal4, UAS-mcherry*, however the loss of *TH* or *Ddc* using this driver did not lead to any changes in Tep4 or P1 areas (Supplementary Fig. 2K). The lack of any P1 expression in *Ddc*^*RNAi*^ condition is because of both the lines being P1 negative (Honti et al., 2013). Taken together, these data implied a significant requirement for intracellular dopamine synthesis in blood progenitor cells for lymph gland growth. The data also demonstrated a requirement for dopamine synthesis in differentiation of blood progenitor cells however the differences in phenotypes detected using *Dome-Gal4* versus *Tep4-Gal4*, suggested its requirement either in the subset of Dome^+^Tep4^-^ progenitor cells or a temporal function of dopamine in progenitor differentiation.

### Extracellular sensing of dopamine via dopamine receptor *Dop2R* and dopamine transporter *(DAT)* in the progenitor cells

Next, we examined the mechanisms by which blood progenitor cells sensed dopamine. Dopamine, can be sensed by dopamine receptors and via the dopamine transporter (DAT) that transports dopamine into cells.

To test if lymph gland blood progenitor cells were responsive to dopamine-based receptor activation, we employed the use of a GPCR activation-based Dopamine (GRAB_DA_) sensor. This fluorescent based sensor enables a sensitive, direct and cell specific detection of extracellular dopamine. The sensor is a dopamine receptor that has been engineered to contain a GFP reporter that gives fluorescence upon dopamine binding induced conformational change in the receptor (Sun et al., 2018). We expressed this sensor (*UAS-DA1m*) in lymph gland progenitor cells using *domeMESO-Gal4* driver. Compared to age matched control lymph glands, where expression of the driver, *domeMESO-Gal4* driving *mCD8::GFP* (Fig. 3A-C), was detected in 70% of cells, the dopamine receptor-based activity sensor was detected only in a subset of these (Fig. 3A’-C’). This localized activity-based fluorescence of the sensor implied dopamine receptor signalling in only specific subset of the blood progenitor cells. The restriction in activity-based fluorescence to a subset is not because of the inability of the construct to be expressed in other cells as when driver line expression was assessed by analysing its ability to drive *UAS-mCD8::GFP* ,(Fig. 3A-C), the expression of GFP was detected in a large population of cells. Compared to this, the expression of the dopamine receptor activity reporter was detected in substantially fewer cells. We also found that as lymph gland development progressed, the number of cells expressing the receptor activity increased over time (Fig. 3A’-C’). At earlier time points, dopamine receptor activity was detected in a small subset of progenitor cells that were localized at the periphery of the lymph gland (Fig. 3A’-B’). As development progressed, cells in the leading edge of the MZ, closer to the CZ (Fig. 3A’-B’) were positive for dopamine receptor activity. By mid 3^rd^ instar this pattern emerged more clearly and the cells at the juxtaposition of MZ and peripheral CZ cells showed active receptor signalling (Fig. 3C’). This pattern of activity was reminiscent of the positioning of the intermediate or transitioning progenitors (Blanco-Obregon et al., 2020) and the sensor expression analysis highlighted dopamine signalling at this interface.

**Fig 3:**
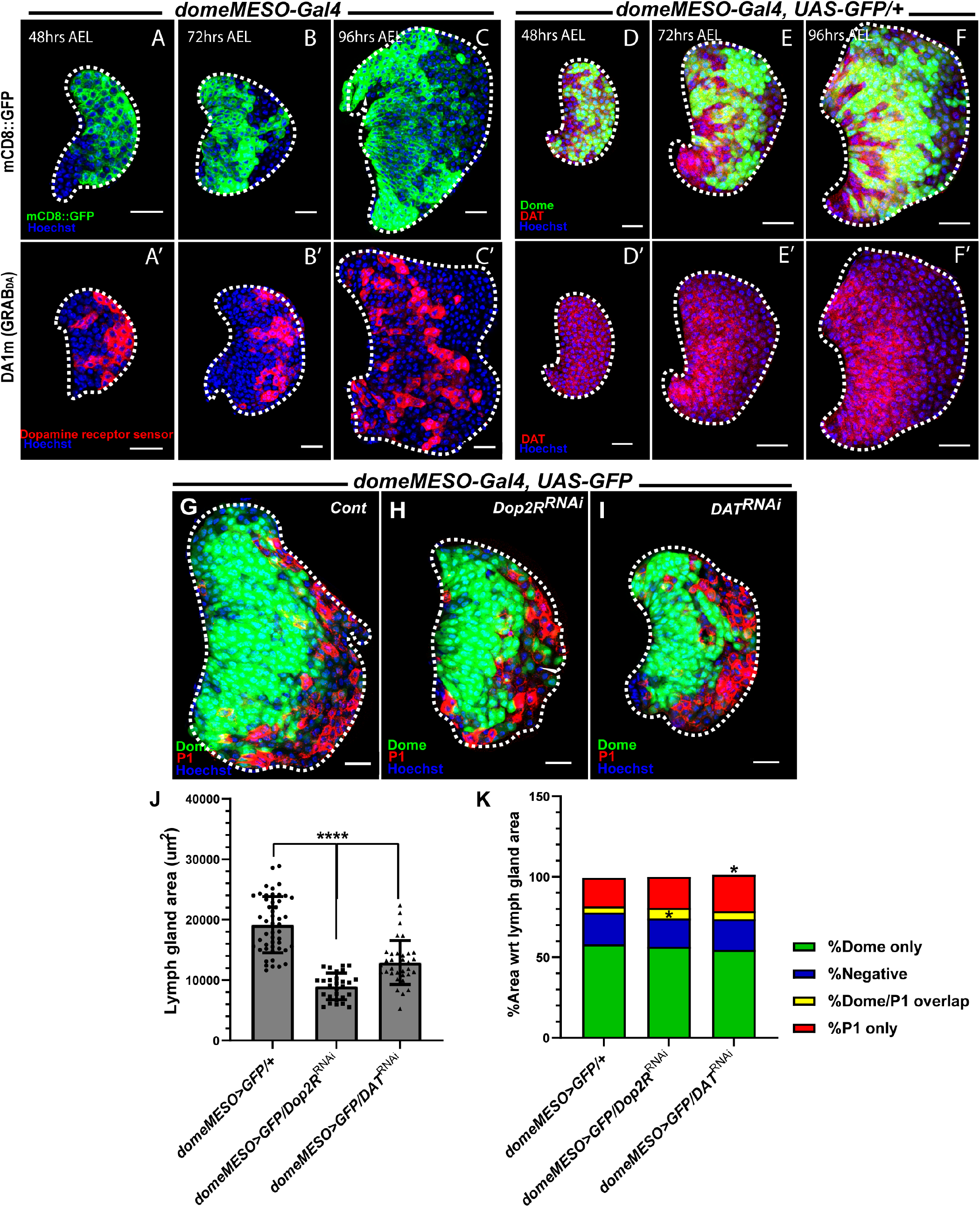
Dopamine sensing in the progenitors via dopamine receptor *Dop2R* and dopamine transporter *DAT* contributes to lymph gland size regulation. DNA is stained with Hoechst in blue, Dopamine transporter (DAT) in red, Dopamine receptor-based reporter (DA1m) in red, *domeMESO* (representative of progenitors) expression in green, P1 (representative of mature plasmatocytes) in red, *mCD8::GFP* in green. All image panels show a 20µm scale bar. AEL indicates <u>A</u>fter <u>E</u>gg <u>L</u>aying. The quantification in J represents mean with standard deviation and in K represents mean. The statistical analysis applied in J and K is unpaired t-test, two tailed. ‘n’ represents the number of lymph gland lobes analyzed. For better representation, the primary lobe of the lymph gland has been represented and outlined in white dashed lines. (A-C’) Expression of dopamine receptor reporter (*UAS-DA1m*), a GPCR based activation reporter, in the progenitors using *domeMESO-Gal4* driver was conducted in a temporal manner-early 2^nd^ (48h AEL, A’), early 3^rd^ (72h AEL, B’) and mid-3^rd^ (96h AEL, C’) instar lymph glands. A subset of the progenitors showed expression of the reporter (active signaling cells) when compared to age matched controls expressing *mCD8::GFP* (A-C). Additionally, an increase in number of cells active for dopamine signaling was found with increasing lymph gland size. (D-F’) Immunostaining with anti-DAT antibody shows transporter expression in the lymph gland primary lobe with higher expression in the inner core region, Dome positive, of the tissue. (G-I) Using *domeMESO-Gal4, UAS-GFP* driver line, *Dop2R* and *DAT* knockdown was conducted and this gave a reduction in lymph size in the (H and I compared to G). (J) Quantification of lymph gland size (*domeMESO-Gal4, UAS-GF/+*, n= 49, *domeMESO-Gal4, UAS-GFP*; *Dop2R*^*RNAi*^, n= 28, ****p<0.0001, *domeMESO-Gal4, UAS-GFP*; *DAT*^*RNAi*^, n= 33, ****p<0.0001). (K) Quantifications of the proportions of % Dome (green bars for MZ), % negative (blue bar), % overlap (Dome and P1, yellow bars) and % P1 (red bars for CZ) suggested no significant differences in the Dome area between the knockdown of *Dop2R* and *DAT* when compared to the control (*domeMESO-Gal4, UAS-GF/+*, n= 26, *domeMESO-Gal4, UAS-GFP*; *Dop2R*^*RNAi*^, n= 16, p=0.6032, *domeMESO-Gal4, UAS-GFP*; *DAT*^*RNAi*^, n= 22, p=0.191) and the % negative area (*domeMESO-Gal4, UAS-GF/+*, n= 27, *domeMESO-Gal4, UAS-GFP*; *Dop2R*^*RNAi*^, n= 16, p=0.3152, *domeMESO-Gal4, UAS-GFP*; *DAT*^*RNAi*^, n= 24, p=0.6586). A slight but significant increase in %overlap (Dome/P1) was observed upon loss of *Dop2R* in the progenitors, which was not the case with *DAT* knockdown (*domeMESO-Gal4, UAS-GF/+*, n= 27, *domeMESO-Gal4, UAS-GFP*; *Dop2R*^*RNAi*^, n= 16, *p=0.0116, *domeMESO-Gal4, UAS-GFP*; *DAT*^*RNAi*^, n= 24, p=0.156). The %P1 population remained unchanged in *Dop2R* loss of function condition (*domeMESO-Gal4, UAS-GFP*; *Dop2R*^*RNAi*^, n= 16, p=0.5235) and had a mild increased in DAT loss of function condition (*domeMESO-Gal4, UAS-GFP*; *DAT*^*RNAi*^, n= 24, *p=0.0314) when compared to control (*domeMESO-Gal4, UAS-GF/+*, n= 27).

Intracellular dopamine availability is also regulated by transport (uptake) through the dopamine transporter, DAT. DAT belongs to the SLC6 family of transporters that couples the Na+/Cl-gradient with the inward movement of dopamine (Vaughan and Foster, 2013). This module regulates the amount of dopamine entering cells from the external milieu. An anti-DAT antibody (Ueno and Kume, 2014) was used to assess DAT expression during lymph gland development. We found substantial levels of DAT throughout cells of the lymph gland (Fig. 3D-F’). Specifically, the expression of DAT was found to be higher in the MZ region (Supplementary Fig. 3_1A-C) of later 3^rd^ instar lymph glands. This pattern was consistent with increasing levels of dopamine (Fig.1), dopamine synthesising enzyme, *Ddc* (Fig. 2G) and receptor activity reporter (Fig. 3C’). These observations suggested that the elevated intracellular dopamine detected in progenitor cells was most likely a consequence of increased synthesis and intracellular transport.

### Extracellular sensing of dopamine via dopamine receptor *Dop2R* and dopamine transporter *(DAT)* in the progenitor cells maintains lymph gland size

This expression pattern of the receptor-activity sensor and transporter led us to investigate the impact of their modulation in progenitor development as undertaken for *TH* and *Ddc. Drosophila* is known to have three kinds of dopamine receptors which are G protein coupled receptors (GPCRs) that includes i) D1 like which includes *Dop1R1* and *Dop1R2* ii) a D2 like that includes *Dop2R* and iii) a non-canonical receptor *DopEcR* that can bind to both ecdysone and dopamine as ligands (Yamamoto and Seto, 2014, Karam et al., 2019). These receptors differ in their mechanism of action downstream by either activating (D1 class and *DopEcR*) or inhibiting (D2 like) adenylate cyclase. In order to address which dopaminergic receptors were of relevance to lymph gland development, we examined the effects of down-regulating each of these in progenitor cells, by driving RNA*i* using *domeMESO-Gal4, UAS-GFP* as the driver. The specific loss of *Dop2R* in the progenitors caused a severe reduction in lymph gland size and recapitulated the *TH* and *Ddc* small lymph gland phenotype (Fig. 3G, H, quantification in J). Unlike synthesis, *Dop2R* loss-of-function condition did not alter the proportions of MZ (marked with GFP) and CZ (marked with P1 in red) which remained unchanged (Fig. 3K). Crystal cell status as assessed by PPO staining also showed no changes (Supplementary Fig. 3_1D, E and quantification in G). This implied that the MZ progenitors of the lymph gland relied on *Dop2R* mediated dopamine sensing to regulate lymph gland size without altering the progenitor/differentiation balance. The loss of other dopamine receptors-*Dop1R1* and *DopEcR* did not result in an appreciable reduction in lymph gland size (Supplementary Fig. 3_1H-J, quantification in L). Lymph gland size dependence on *Dop2R* was further strengthened by i) the analysis using another RNA*i* line for *Dop2R* (BDSC 26001) (Supplementary Fig. 3_1K and quantification in L) ii) as well as using another MZ specific driver, *Tep4-Gal4, UAS-mCherry* (Supplementary Fig. 3_1M, N and quantification in P). In both conditions, a small lymph gland size was recapitulated with no difference in proportions of Tep4 and P1 positive areas (Supplementary Fig. 3_1Q).

The down-regulation of *DAT* in progenitor cells also resulted in small lymph glands (Fig. 3G, I and quantification in J) with no defects in the maintenance and differentiation of progenitor cells, as detected by %P1 area (Fig. 3K). Employing another MZ specific Gal4 driver, *Tep4-Gal4, UAS-mCherry*, to express *DAT*^*RNAi*^ recapitulated the small lymph gland size phenotype (Supplementary Fig.3_1M, O and quantifications in P) and this also did not lead to any defect of Tep4 and P1 areas (Supplementary Fig.3_1M, O and quantifications in Q). Assessment of crystal cells did not show any defects upon progenitor loss of *DAT* using *domeMESO-Gal4, UAS-GFP* driver (Supplementary Fig. 3_1D, F and quantification in G). These findings showed that unlike dopamine synthesis, extracellular dopamine sensing via *Dop2R* or uptake via *DAT* in the progenitor cells allowed lymph gland growth without perturbing their maintenance or differentiation status.

### Lymph gland growth kinetics

The small lymph gland size phenotype was a common observation in all dopamine modules; synthesis, receptor-based sensing via *Dop2R* and uptake via the transporter. Importantly, their down-regulation in the differentiating blood cells using *Hml*^*Δ*^*-Gal4, UAS-2xEGFP* did not result in any change in the sizes of the tissue (Supplementary Fig. 3_2A-F). These results suggested that dopamine is predominantly required by the lymph gland blood progenitor cells to coordinate lymph gland growth.

To assess the underlying reasons for the small size of the lymph gland, we investigated if any increase in cell death would account for the phenotype. We used cleaved-caspase 3 staining to mark apoptotic nuclei (Grigorian et al., 2013) in late 3^rd^ instar lymph glands, genetically manipulated for loss of either dopamine synthesis or sensing proteins in progenitor cells. As a positive control, *Hml*^*Δ*^*-Gal4, UAS-GFP, UAS-Hid* expressing lymph glands were used, wherein apoptosis was induced and caspase 3 puncta around the nucleus were detected (Supplementary Fig. 4_1A-A’, inset in A’’). Here, we failed to detect any apoptotic nuclei (Supplementary Fig. 4_1B-F’’) in these genetic conditions. This showed that the size reduction was not a consequence of increased cellular death.

**Fig. 4:**
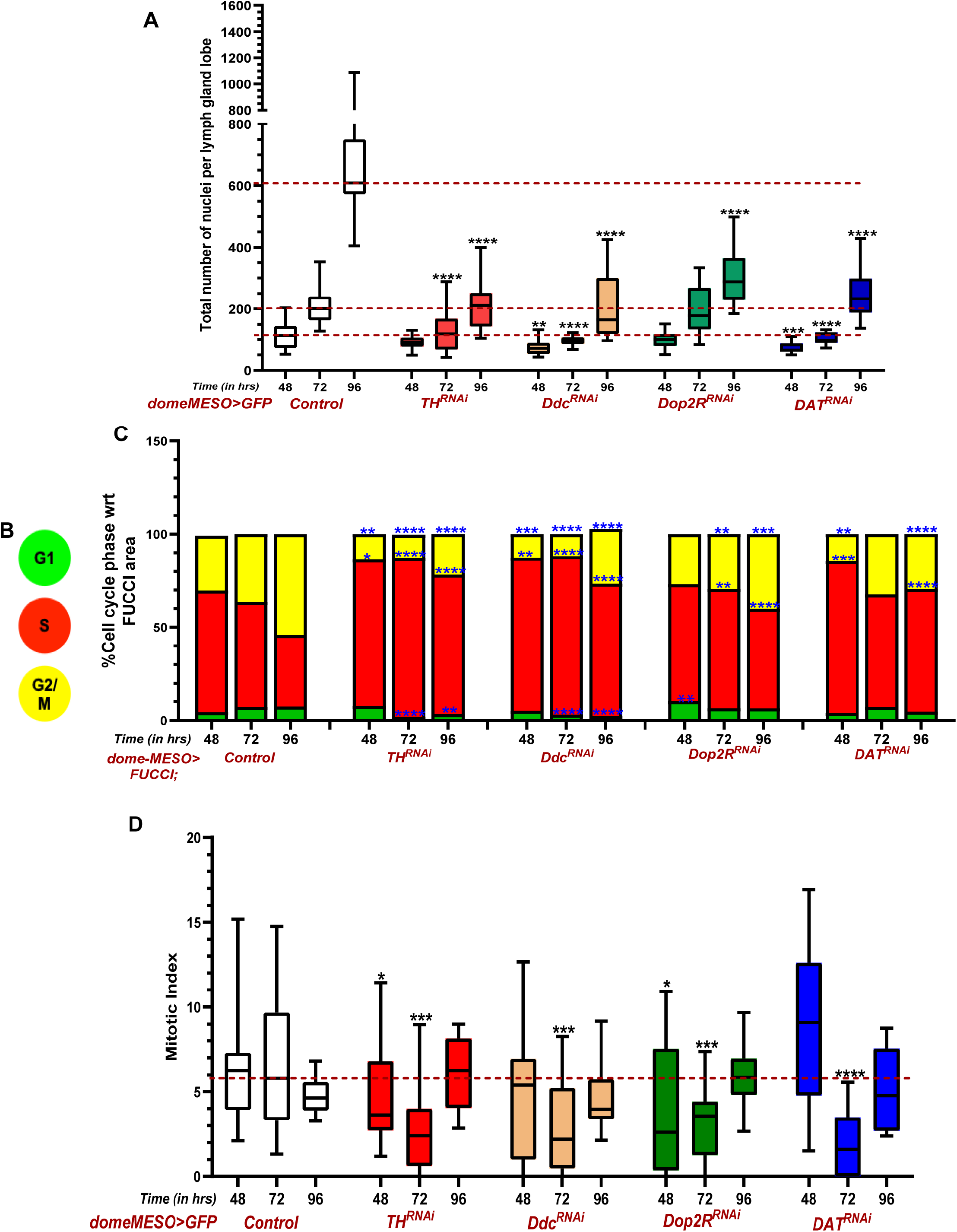
The trilogy of dopamine sensing, synthesis and uptake regulate lymph gland growth and show differential effects on cell cycle phasing. DNA is stained with Hoechst in blue, FUCCI (fluorescent ubiquitination-based cell cycle indicator) reporter shows G1 phase cells in green, S phase cells in red and G2/M phase cells in yellow. All panels show a 20µm scale bar. AEL indicates <u>A</u>fter <u>E</u>gg <u>L</u>aying. The quantification in A represents median with box plots and that in R represents mean. The statistical analysis applied in A and R is Mann-Whitney test while. ‘n’ represents the number of lymph gland lobes analyzed. For better representation, the primary lobe of the lymph gland has been represented and outlined in white dashed lines. (A) An assessment of total nuclei counts in a temporal manner was conducted upon perturbation of dopamine modules of synthesis, signaling via *Dop2R* and uptake through *DAT* in the progenitors using *domeMESO-Gal4, UAS-GFP* driver. These perturbations resulted in reduction of total nuclei counts upon downregulation of *TH, Ddc* and *DAT* at 48h AEL (*domeMESO-Gal4, UAS-GF/+*, n= 36, *domeMESO-Gal4, UAS-GFP*; *TH*^*RNAi*^, n= 21, p=0.0718, *domeMESO-Gal4, UAS-GFP*; *Ddc*^*RNAi*^, n= 16, **p=0.0016, *domeMESO-Gal4, UAS-GFP*; *DAT*^*RNAi*^, n= 15, ***p=0.0008). The loss of *Dop2R* did not show any significant reduction at this time (*domeMESO-Gal4, UAS-GFP*; *Dop2R*^*RNAi*^, n= 21, p=0.2374). Later, at 72h AEL, the total number of nuclei remained significantly low in *TH, Ddc* and *DAT* knockdowns (*domeMESO-Gal4, UAS-GF/+*, n= 26, *domeMESO-Gal4, UAS-GFP*; *TH*^*RNAi*^, n= 21, ****p<0.0001, *domeMESO-Gal4, UAS-GFP*; *Ddc*^*RNAi*^, n= 17, ****p<0.0001, *domeMESO-Gal4, UAS-GFP*; *DAT*^*RNAi*^, n= 22, ****p<0.0001). This was still not the case for *Dop2R* loss of function condition (*domeMESO-Gal4, UAS-GFP*; *Dop2R*^*RNAi*^, n= 18, p=0.126). At 96h AEL, the perturbation of all these modules reflected a reduction in total nuclei counts (*domeMESO-Gal4, UAS-GF/+*, n= 18, *domeMESO-Gal4, UAS-GFP*; *TH*^*RNAi*^, n= 21, ****p<0.0001, *domeMESO-Gal4, UAS-GFP*; *Ddc*^*RNAi*^, n= 17, ****p<0.0001, *domeMESO-Gal4, UAS-GFP*; *DAT*^*RNAi*^, n= 22, ****p<0.0001, *domeMESO-Gal4, UAS-GFP*; *Dop2R*^*RNAi*^, n= 20, ****p<0.0001). (B) Color coding scheme to represent the cell cycle phase as shown through FUCCI reporter. (C) Quantifications of the % cell cycle phase i.e., %G1, %S and %G2/M at different time points, in the dopamine perturbed backgrounds. For 48h AEL, significant differences were found in synthesis and uptake module compared to control (*domeMESO-Gal4; UAS-FUCCI/+*, n=34 [%G1], n=35 [%S], n=35 [%G2/M], *domeMESO-Gal4; UAS-FUCCI; TH*^*RNAi*^, n= 13 [%G1, p=0.2215], n=13 [%S, *p=0.0312], n=13 [%G2/M, **p=0.0023], *domeMESO-Gal4; UAS-FUCCI; Ddc*^*RNAi*^, n= 15 [%G1, p=0.9865], n=15 [%S, **p=0.0014], n=15 [%G2/M, ***p=0.0004], *domeMESO-Gal4; UAS-FUCCI; DAT*^*RNAi*^, n= 17 [%G1, p=0.6069], n=17 [%S, ***p=0.001], n=17 [%G2/M, **p=0.0019]). This accumulation in S phase was not found in *Dop2R* loss of function condition, although an increase in G1 phase was noticed (*domeMESO-Gal4; UAS-FUCCI; Dop2R*^*RNAi*^, n= 19 [%G1, **p=0.0013], n=19 [%S, p=0.7569], n=19 [%G2/M, p=0.453]). At 72h, *Dop2R* loss of function started showing an S phase accumulation with reduction in G2/M along with loss of synthesis modules which however was not demonstrated by *DAT* perturbation (*domeMESO-Gal4; UAS-FUCCI/+*, n=38 [%G1], n=38 [%S], n=38 [%G2/M], *domeMESO-Gal4; UAS-FUCCI; TH*^*RNAi*^, n= 22 [%G1, ****p<0.0001], n=19 [%S, ****p<0.0001], n=15 [%G2/M, ****p<0.0001], *domeMESO-Gal4; UAS-FUCCI; Ddc*^*RNAi*^, n= 29 [%G1, ****p<0.0001], n=26 [%S, ****p<0.0001], n=22 [%G2/M, ****p<0.0001], *domeMESO-Gal4; UAS-FUCCI; Dop2R*^*RNAi*^, n= 47 [%G1, p=0.4194], n=47 [%S, **p=0.0047], n=47 [%G2/M, **p=0.0042], *domeMESO-Gal4; UAS-FUCCI; DAT*^*RNAi*^, n= 29 [%G1, p=0.7557], n=29 [%S, p=0.2322], n=29 [%G2/M, p=0.1863]). At 96h AEL, all the modules showed an expansion of S phase with decrease in G2/M phase, albeit with differences in the levels (*domeMESO-Gal4; UAS-FUCCI/+*, n=55 [%G1], n=55 [%S], n=55 [%G2/M], *domeMESO-Gal4; UAS-FUCCI; TH*^*RNAi*^, n= 14 [%G1, **p=0.0047], n=14 [%S, ****p<0.0001], n=14 [%G2/M, ****p<0.0001], *domeMESO-Gal4; UAS-FUCCI; Ddc*^*RNAi*^, n= 20 [%G1, ****p<0.0001], n=19 [%S, ****p<0.0001], n=19 [%G2/M, ****p<0.0001], *domeMESO-Gal4; UAS-FUCCI; Dop2R*^*RNAi*^, n= 23 [%G1, p=0.9761], n=23 [%S, ****p<0.0001], n=23 [%G2/M, ***p=0.0004], *domeMESO-Gal4; UAS-FUCCI; DAT*^*RNAi*^, n= 19 [%G1, p=0.0872], n=19 [%S, ****p<0.0001], n=19 [%G2/M, ****p<0.0001]). (D) Quantification of the mitotic index in a time course manner with dopamine modules perturbation in the progenitors using *domeMESO-Gal4; UAS-GFP* driver. At 48h, mild reduction was observed in *TH* and *Dop2R* loss of function condition (*domeMESO-Gal4; UAS-GFP/+*, n=33, *domeMESO-Gal4; UAS-GFP; TH*^*RNAi*^, n=18, *p=0.0498, *domeMESO-Gal4; UAS-GFP; Ddc*^*RNAi*^, n=16, p=0.4187, *domeMESO-Gal4; UAS-GFP; Dop2R*^*RNAi*^, n=21, *p=0.0219, *domeMESO-Gal4; UAS-GFP; DAT*^*RNAi*^, n=15, p=0.1215). At 72h, a significant reduction in the mitotic index was observed (*domeMESO-Gal4; UAS-GFP/+*, n=28, *domeMESO-Gal4; UAS-GFP; TH*^*RNAi*^, n=23, ***p=0.0002, *domeMESO-Gal4; UAS-GFP; Ddc*^*RNAi*^, n=17, ***p=0.001, *domeMESO-Gal4; UAS-GFP; Dop2R*^*RNAi*^, n=25, ***p=0.0003, *domeMESO-Gal4; UAS-GFP; DAT*^*RNAi*^, n=27, ****p<0.0001). At 96h, no significant reduction was found (*domeMESO-Gal4; UAS-GFP/+*, n=16, *domeMESO-Gal4; UAS-GFP; TH*^*RNAi*^, n=21, p=0.0726, *domeMESO-Gal4; UAS-GFP; Ddc*^*RNAi*^, n=16, p=0.6088, *domeMESO-Gal4; UAS-GFP; Dop2R*^*RNAi*^, n=18, *p=0.0271, *domeMESO-Gal4; UAS-GFP; DAT*^*RNAi*^, n=21, p=0.5144).

The lymph gland growth is accomplished by rapid proliferation of progenitor cells at earlier stages of development which gradually ceases later on (Jung et al., 2005, Krzemien et al., 2010, Mondal et al., 2011, Sharma et al., 2019). Hence, we next undertook a comparative analysis of growth profiles of control and mutant lymph glands lacking dopamine function in progenitor cells. We estimated total cell counts of lymph gland primary lobes (please see Methods) that were obtained from larvae at 48, 72 and 96h AEL for this analysis.

We observed that in control conditions, lymph gland cell numbers doubled from 48 to 72h AEL and tripled from 72 to 96h AEL (Fig. 4A). The late proliferative burst occurring at 72h was unexpected and has not been described previously. In conditions lacking dopamine function in the progenitors, either synthesis (*TH* and *Ddc* RNAi) or sensing (*Dop2R* and *DAT* RNAi), we observed an overall reduction in total cell numbers at 96h AEL (Fig. 4A). This confirmed that the lymph gland size reduction was a consequence of fewer number of cells. When assessed at an earlier developmental stage at 48h AEL, in all conditions genetically manipulated for dopamine, the total cell numbers were either mildly reduced or comparable to controls (Fig. 4A). At 72h AEL, when the control had doubled its cell numbers, the synthesis and transporter mutants failed to demonstrate any increase and the total cell numbers remained comparable to cell counts detected at 48h AEL (Fig. 4A, please see the brown dashed line for control’s reference at each time point). This implied that after 48h, dopamine synthesis and uptake was necessary to build intracellular dopamine to support the doubling of progenitor cell numbers. Contrary to this, loss of dopamine receptor function showed comparable total cell numbers both at 48 and 72h AEL. This implied that despite the loss of progenitor *Dop2R* function, cell counts doubled and lymph glands grew comparable to controls. From 72 to 96h AEL, however, when the controls tripled their cell counts, the *Dop2R* mutants showed a marginal doubling in cell numbers. These data suggested a temporal requirement for *Dop2R* signalling in ramping lymph gland cell numbers from 72 to 96h AEL as opposed to synthesis, which was required from earlier stages of its development.

### Dual control of dopamine in the regulation of lymph gland proliferation

We analysed cell cycle phasing to gain insight into how cell numbers increased during lymph gland development. For this, we employed the use of the FUCCI reporter, a fluorescent ubiquitination-based cell cycle indicator, and performed a comparative assessment of cell cycle profiles. The FUCCI fluorescence assay gives details on stages of cell cycle progression and utilizes two probes that are sensitive to ubiquitination-based degradation by ubiquitin E3 ligases, depending on the state of the cell cycle. The first probe contains a GFP reporter fused to the E2F protein that is targeted by the CRL4^Cdt2^ ubiquitin ligase. This degradation step occurs in the S phase of the cell cycle. The second construct harbours monomeric red fluorescent protein as the reporter tagged to the Cyclin B (regulator of G2 to M phase). This Cyclin B protein is targeted for degradation by APC/C complex in the G1 phase of the cell cycle. Thus, based on the fluorochrome expression, it can be deciphered whether the cells are in G1 (green fluorescence), S (red fluorescence) or G2/M phase (yellow fluorescence) (Zielke *et al*., 2014, Fig. 4B).

A temporal analysis of progenitor cell cycle states respectively at 48, 72 and 96h AEL was undertaken and the percentage cell number in G1, S or G2/M phases of the cell cycle, was assessed. For this, the number of cells in each phase of the cell cycle were counted and represented against the total number of cells positive for the reporter (See Methods for detail). Using this approach, we observed that in control larvae at 48h, when the lymph gland comprises only of Dome^+^ progenitor cells, almost 65% of progenitor cells, were detected in the S phase, with 30% in G2/M and a small population of 5% cells in G1 phase of the cell cycle (Fig. 4C and Supplementary Fig. 4_2A). Another 24 hrs into development, at 72h AEL, a decline in the proportion of S phase cells was noticed. This was accompanied by a rise in the G2/M population and a small increase in G1 cells (Fig. 4C and Supplementary Fig. 4_2F). The cell cycle status at 72h, coincides with progenitor differentiation and the formation of a distinct CZ. By mid-3^rd^ instar, at 96h AEL, a majority of progenitor cells (around 60%) were now in the G2/M phase while the S phase population had substantially reduced to 30-35% (Fig. 4C and Supplementary Fig. 4_2K). Interestingly, the G1 phase was maintained at 5-7% of the progenitor population (Fig. 4C and Supplementary Fig. 4_2A, F and K). This suggested, that progenitor cells earlier in development were maintained in S phase of the cell cycle and gradually transitioned into G2 which ultimately led to majority being retained in G2 by 96h AEL. The accumulation of progenitor cells in G2 was consistent with published data (Sharma et al., 2019) and our data here highlighted the cell cycle trajectory over developmental time course that led to G2.

The FUCCI yellow marker, marks cells both in G2 and M phase (Fig. 4B).Therefore, to gauge the number of cells mitotically active in the yellow/G2 subset, the mitotic index at the different time points was characterized. We undertook staining to mark for phosphorylation of Histone (pH3), which is a robust marker of mitotically active cells, as the serine residues 10 and 28 are heavily phosphorylated in the metaphase stage of the cell cycle (Ye-Kim *et al*., 2017). We undertook pH3 analysis at 48, 72 and 96h AEL (Fig. 5A-O’) and estimated the percentage of mitotically active cells in the lymph gland. We observed that, in controls, throughout development, the mitotic index remained fairly consistent at 5% (Fig. 4D). Which implied that the increasing proportion of progenitor population into G2 during development increased the probability of cells to enter M phase but at any given time point only 5% cells were mitotically active. We speculate that the developmental requirement to maintain higher number of cells in G2 by 72h AEL, increased the overall mitotic potential of the lymph gland. This may be necessary to enable the rapid increase in lymph gland size detected from 72 to 96h AEL (Fig. 4A) and prime their entry into differentiation trajectory (Mondal *et*.*al*., 2011).

**Fig. 5:**
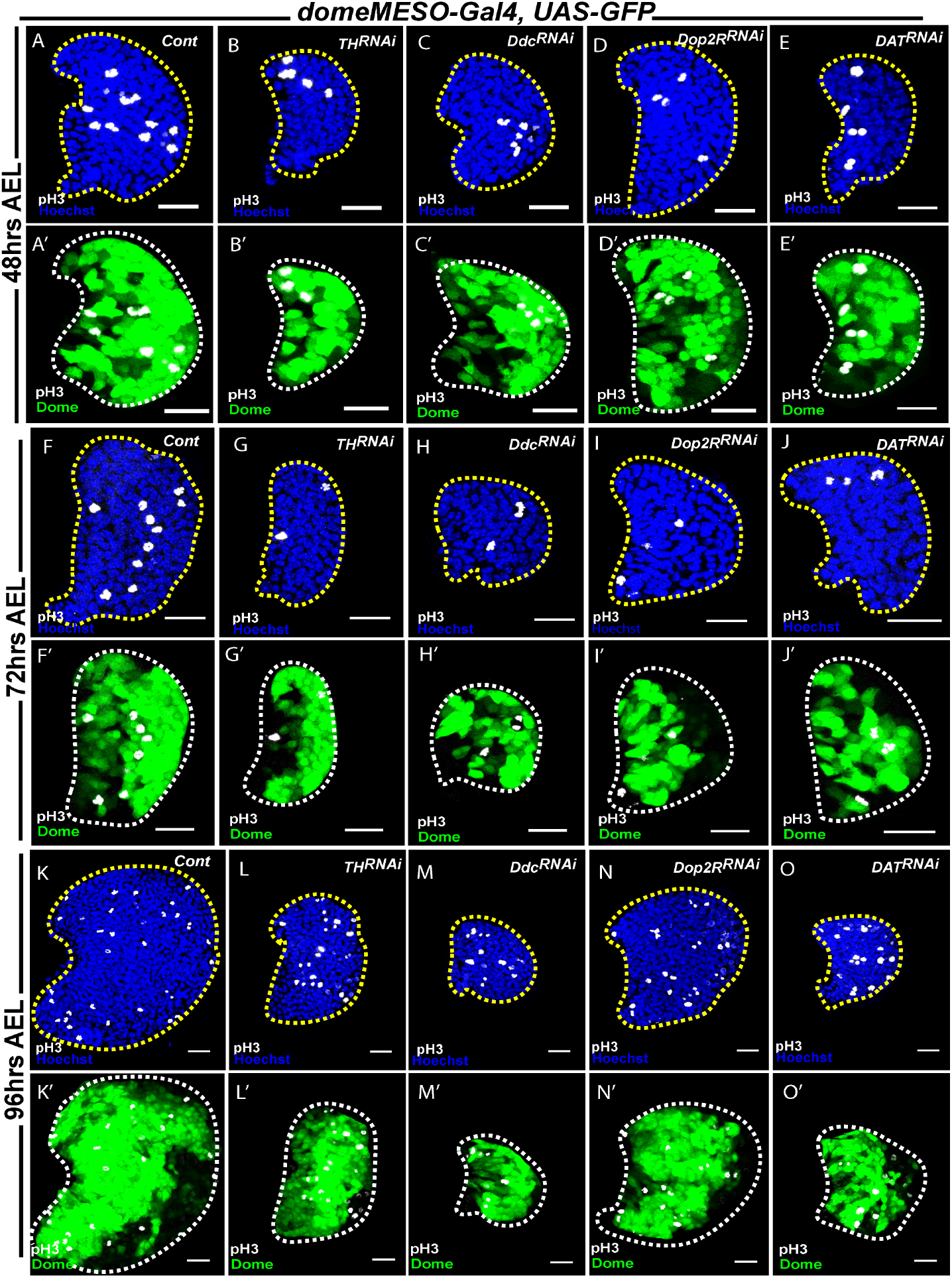
Dopamine exhibits dual modulation on lymph gland growth dynamics. DNA is stained with Hoechst in blue, *domeMESO* (representative of progenitors) in green, pH3 (phospho-Histone 3) in grey. All panels show a 20µm scale bar. AEL indicates <u>A</u>fter <u>E</u>gg <u>L</u>aying. The quantification in P represents median with box plots. The statistical analysis applied in P is Mann-Whitney test. ‘n’ represents the number of lymph gland lobes analyzed. For better representation, the primary lobe of the lymph gland has been represented and outlined in white dashed lines. (A-E’) Representative lymph gland images at 48h AEL reflecting proliferation profile (via pH3 analysis) in control (A-A’) and loss of function of function of *TH* (B-B’), *Ddc* (C-C’), *Dop2R* (D-D’) and *DAT* (E-E’) using *domeMESO-Gal4; UAS-GFP* driver. No significant differences in mitotic index were observed in the dopamine perturbed backgrounds when compared to control. (F-J’) Representative lymph gland images at 72h AEL reflecting proliferation profile (via pH3 analysis) in control (F-F’) and loss of function of function of *TH* (G-G’), *Ddc* (H-H’), *Dop2R* (I-I’) and *DAT* (J-J’) using *domeMESO-Gal4; UAS-GFP* driver. An overall reduction in mitotic index was observed in the dopamine perturbed backgrounds when compared to control. (K-O’) Representative lymph gland images at 96h AEL reflecting proliferation profile (via pH3 analysis) in control (K-K’) and loss of function of function of *TH* (L-L’), *Ddc* (M-M’), *Dop2R* (N-N’) and *DAT* (O-O’) using *domeMESO-Gal4; UAS-GFP* driver. Here *TH* and *Dop2R* showed milder reduction in mitotic index when compared to control which was not the case for *Ddc* and *DAT* loss of function condition.

The FUCCI tool was next driven in progenitor cells lacking dopamine synthesis, signalling and uptake in a temporal manner (Supplementary Fig. 4_2A-O) and analysed as above. Upon progenitor specific loss of dopamine synthesis via *TH* or *Ddc*, the FUCCI analysis revealed that at 48h AEL, a significant expansion in the S phase population was evident (Supplementary Fig. 4_2B-C compared to A). About 80-85% cells were detected in S phase of their cell cycle with only 12-14% in G2/M (Fig. 4C). This cell cycle state was consistently seen over time at 72 and 96h AEL (Supplementary Fig. 4_2G-H compared to F and L-M compared to K, quantification in Fig. 4C). Thus, a reduction in total cell counts seen in these mutants (Fig. 4A) and the accompanying growth defect was a consequence of cells being maintained in S phase of the cell cycle. These data implied a function for intracellular dopamine synthesis in driving exit of cells from S phase of the cell cycle.

Loss of *DAT* function recapitulated a similar increase in S phase cells at 48h AEL with a reduction in G2/M phase (Supplementary Fig. 4_2E compared to A, quantification in Fig. 4C). At 72 and 96h AEL, the progenitors with loss of *DAT* function showed lesser proportion in S phase with an increase in G2/M phase, however the number of cells in S phase continued to remain moderately higher than seen in controls (Supplementary Fig. 4_2J compared to F, O compared to K, quantification in Fig. 4C). The loss of *DAT* data corroborated with a requirement for intracellular dopamine as seen with *TH* and *Ddc* and suggested that in blood progenitor, the gradual increase in intracellular dopamine level was a consequence of both synthesis and increased *DAT* mediated uptake. This may be necessary for progenitor cells to reach a certain threshold of intracellular dopamine required to exit S and enter G2. The mitotic index at 48 and 72h AEL, upon loss of dopamine synthesis (Fig. 5B-C’ compared to A-A’ for 48h and G-H’ compared to F-F’ for 72h) or transporter (Fig. 5E-E’ for 48h and J-J’ for 72h) showed a significant reduction (Fig. 4D), which was consistent with the increased S and reduced G2 phase cells detected in these conditions (Fig. 4B). This explained the reduction in the total cell counts observed in the mutant lymph gland (Fig. 4A). At 96h, the mitotic index however did not show any difference and was restored back to control levels (Fig. 5L-M’ for synthesis loss of function and O-O’ for *DAT* loss of function, quantification in Fig. 4D) which suggested dopamine independent control of lymph gland development at later stages.

Loss of dopamine signalling via *Dop2R*, however showed no difference at 48h in terms of S, G2/M cell populations (with a minor increase in G1) and was unlike loss of dopamine synthesis or uptake (*TH* or *Ddc* or *DAT* knockdown) (Supplementary Fig. 4_2D compared to A, quantification in Fig. 4C). Later, at 72 or 96h only minor increase in S and minor reduction in G2/M was noticeable (Supplementary Fig. 4_2I compared to F for 72h, N compared to K for 96h, quantification in Fig. 4C). This implied that the reduction in total cell numbers seen from 72 to 96h AEL (Fig. 4A) was not a consequence of any alteration in S or G2 cell cycle phasing. We assessed the mitotic index at these time points and observed a significant reduction specifically at 72h. (Fig. 5I-I’ compared to F-F’, quantification in Fig. 4D). At 48h, or later at 96h, the mitotic potential remained comparable to control, which implied that the mitotic potential at 72h (Fig. 4A) was under the control of dopamine receptor signaling. After this time point, the mitotic index was independent of dopamine regulation (Fig. 5N-N’ compared to K-K’, quantification in Fig. 4D). Based on these data we propose that, at 72h AEL, the lymph gland progenitor cells lacking dopamine receptor signalling had reduced mitosis (Supplementary Fig. 5A), hence the number of total cells ended up being lesser at 96h (Fig. 4A). These data revealed a temporal requirement for *Dop2R* function from 72h onwards until 96h for driving the proliferative boost seen in control lymph glands. Consistent with this result, cells positive for dopamine receptor sensor were positive for pH3 only at 72 h and not at earlier time point (Supplementary Fig. 5B-D, yellow arrows for indicating pH3 positive either non-overlapping with the sensor (B) or overlapping (C, D), see inset in panel C).

Previous studies have implicated the use of neurotransmitters by progenitor cells in their maintenance (Shim et al., 2013) and immune response (Madhwal et al., 2020) and by mature immune cells for their function (Qi et al., 2018). These findings led us to explore other neurotransmitters in blood development and our work in this study features dopamine in controlling a fundamental process of lymph gland growth. The current study showcases the utilization of dopamine by progenitor cells as an essential proliferative cue to coordinate their cell cycle progression and achieve growth of the hematopoietic tissue.

Dopamine, a member of the catecholamine family, is obtained from tyrosine. Tyrosine hydroxylase (TH) catalyses the rate limiting conversion of tyrosine to the intermediate L-Dopa, which is converted to dopamine through Dopa decarboxylase (Ddc) dependent reaction. Dopamine is packed in vesicles via vesicular monoamine transporter (VMATs) to be subsequently released in the synapse enabling its recycling. The receiving cells sense dopamine by either D1 or D2 class of receptors (in vertebrates) (Yamamoto and Vernier, 2011) or *DopR, Dop2R* and *DopEcR* (in *Drosophila*) (Yamamoto and Seto, 2014). Dopamine transporter (*DAT*) is known to re-uptake dopamine from the synaptic cleft into the presynaptic neuron once the downstream postsynaptic neuron has received the stimulus (Vaughan and Foster, 2013). As a neurotransmitter, dopamine is well recognised for its various roles in arousal (Kume et al., 2005), motivation (Wise, 2004), learning and forgetting (Berry et al., 2012) to name a few. It’s function in enabling proliferation of adult neural precursor cell (O’Keeffe C et al., 2009) and immature human CD34^+^ hematopoietic progenitor cells (Spiegel et al., 2007) are findings which exhibits growth regulatory aspect of this neurotransmitter, but mainly focused through the lens of receptor mediated signalling.

Our data proposes a non-neuronal function of dopamine in blood progenitor cells that is necessary to moderate the overall growth of the lymph gland (Fig. 6). Growth of this blood forming tissue is supported by dual function of dopamine. Initially, as the lymph gland begins its growth, progenitors are mainly in S phase of the cell cycle. Dopamine synthesis is initiated in early progenitor cells and a threshold of intracellular dopamine is built over time, by utilizing both synthesis and uptake (Fig. 6, shown by red triangle). The increased availability of intracellular dopamine in progenitor cells, facilitates their exit from S and promotes entry into G2. The increased accumulation in G2 by 72h is also the time when the lymph gland begins to rapidly grow in size. At this time, an increase in *Dop2R* activity in progenitor cells primes their potential to exit G2. We hypothesize, that the entry into mitosis is governed by an additional “input”. We propose that cells that are in G2 are activated for *Dop2R* signalling and are competent to enter mitosis, such that in the presence of this additional trigger, the cells readily enter mitosis. Thus, with time as more *Dop2R* activated cells become available, the mitotic capacity of progenitor cells is increased. Signalling by molecules like adenosine or GABA which have been implicated in lymph gland growth and proliferation are potential candidates (Mondal et al., 2011, Shim et al., 2013). This allows the rapid increase in cell numbers. The mechanisms by which dopamine intracellularly manifests exit of cells from the S phase or entry into G2 remains unclear. In this regard, either processes linked to dopamine breakdown or secondary modifications, as described for serotonin (Walther et al., 2011), dopaminylation (Lepack et al., 2020) of proteins can be speculated as a possibility, but this remains to be tested.

**Fig. 6:**
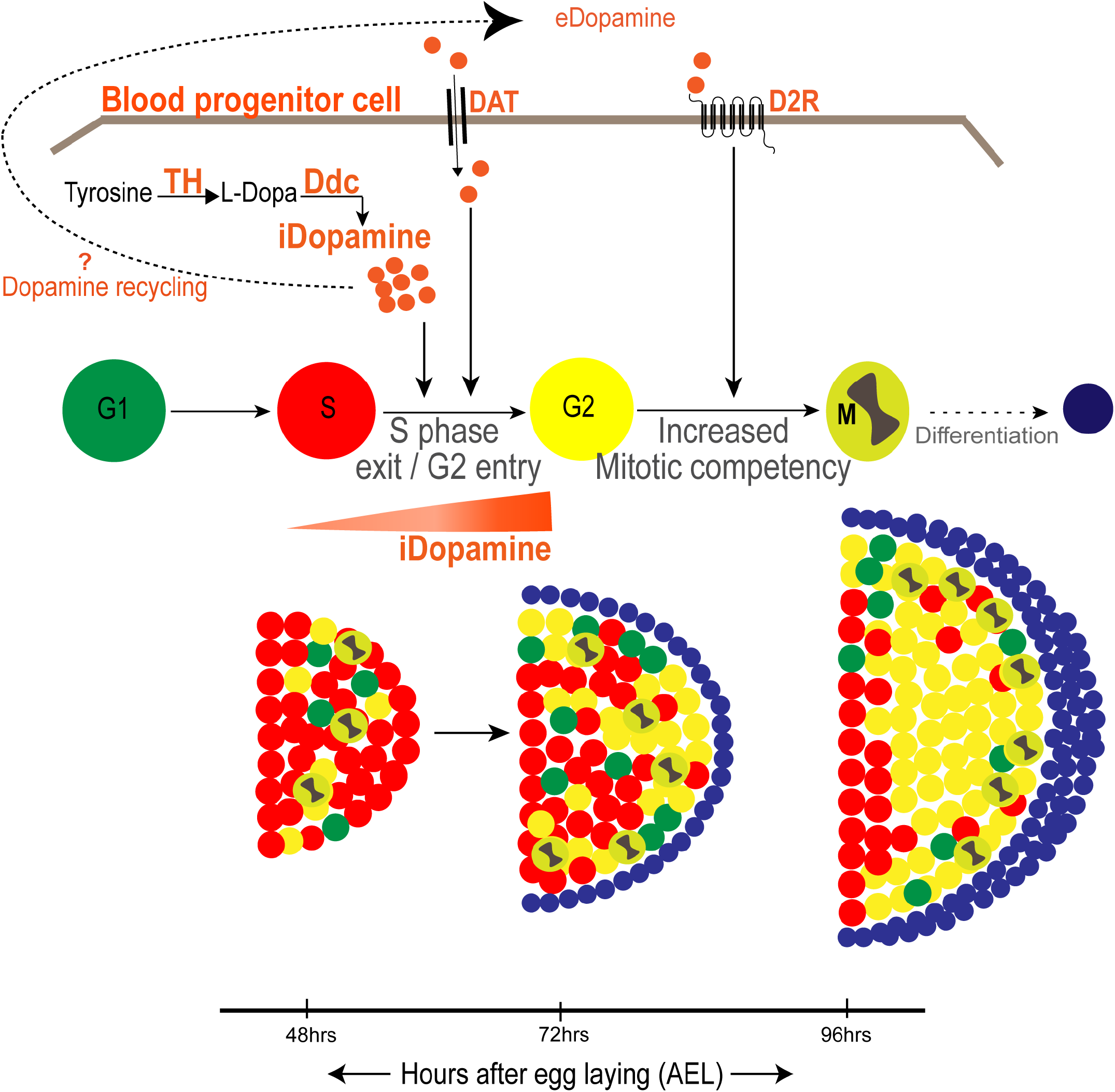
Non-canonical function of dopamine, a neurotransmitter, in enabling progenitor proliferation to achieve lymph gland growth. The study presented here highlights the non-canonical function of dopamine in the proliferation of the hematopoietic progenitor population. The progenitor cells employ the module of dopamine synthesis (enzymes *TH* and *Ddc*) and transporter (*DAT*) to build up the levels of intracellular dopamine (iDopamine). During the earlier stages of lymph gland development, specifically at 48 to 72h AEL, this reservoir of dopamine (as depicted by the red triangle) enables the progenitor pool to transit from S to G2 phase of the cell cycle, Further, the *Dop2R* mediated dopamine signaling, that occurs in a subset of the progenitor population, confers mitotic competency to these progenitor cells enabling them to transit from G2 to M phase of the cell cycle. We also speculate the contributions of dopamine synthesis and uptake in this regard. This culminates in the lymph gland growth to an appropriate size. Overall, dopamine functions not only as a signaling entity but also as an intracellular metabolite to regulate proliferation and cell cycle phasing.

Apart from a few studies which have described dopamine’s role in early neurogenesis (Höglinger et al., 2004, O’Keeffe C et al., 2009), the importance of dopamine as a proliferative cue is not established. Dopamine’s utilization by blood cells as a pro-growth and a mitotic cue both at the level of it being a metabolite and as a signalling molecule are significant findings that illustrate its non-canonical functions in blood development. Our work presented here broadens the understanding of neurotransmitters and their function in non-canonical contexts. The canonical attribution of neurotransmitters in the regulation of brain functions along with neuronal innervations to different tissues where their influence is exerted is most well established. The importance of intracellular dopamine synthesis as a growth signal in blood progenitor cells is unexpected and opens avenues for investigating dopamine from this perspective, which has not yet been undertaken both in neuronal development and also in the hematopoietic field.

## Materials and Method

### *Drosophila* husbandry, stocks, genetics

The following *Drosophila* stocks were used in this study: *yw; +/+; +/+* (wild type, *wt*) *domeMESO-Gal4, UAS-EYFP* (U. Banerjee), *Hml*^*Δ*^*-Gal4, UAS-2xEGFP* (S. Sinenko), *Tep4-Gal4, UAS-mCherry* (U. Banerjee), *domeMESO-Gal4* (U. Banerjee), *Tep4-Gal4* (L. Mandal). The RNAi stocks were obtained from Bloomington (BL) *Drosophila* stock centres. The lines used for the study are: *Dop1R1*^*RNAi*^ (BL 62193), *Dop2R*^*RNAi*^ (BL 26001, Neckameyer and Bhatt, 2012, Inagaki et al., 2012, BL 36824), *DopEcR*^*RNAi*^ (BL 31981, Aranda et al., 2017), *DDC*^*RNAi*^ (BL 27030, Smith and Macdonald, 2020), *TH*^*RNAi*^ (BL 25796, Vaccaro et al., 2017), *DAT*^*RNAi*^ (BL 50619, Jakšic’ et al., 2020), *UAS-FUCCI* (BL 55121), *UAS-DA1m* (BL 80047), *UAS-Hid* (BL 65403), *UAS-mCD8::GFP (*BDSC 5137). All fly stocks were reared on standard corn meal agar food medium with yeast supplementation at 25°C incubator unless specified. The crosses involving RNAi lines were maintained at 29°C to maximize the efficacy of the *Gal4-UAS RNAi* system. The controls in the study correspond to either *yw; +/+; +/+* or Gal4 drivers crossed with *yw; +/+; +/+*.

### Immunostaining and Immunohistochemistry

Lymph glands isolated from larvae were stained following the staining protocol as described previously (Jung et al., 2005). Briefly, the lymph gland tissues from synchronised larvae of a specific developmental stage were dissected in 1X PBS (phosphate buffered saline) from. The tissues were then fixed in 4% formaldehyde for 20 mins followed by three washes with 0.3% PBT (1X PBS + 0.3% Triton X 100) of 10 mins each. Blocking was next carried out in 5% NGS (normal goat serum) for 30-45 mins and the samples were then subjected to primary antibody (Ab) staining of specific dilutions in an overnight, 4°C condition. After the primary antibody treatment, the tissues were washed with 0.3% PBT thrice for 10 mins each to remove the residual antibody. Post this step, a secondary antibody treatment was employed wherein an Alexa Flour conjugated secondary antibody and Hoechst 33342 (for nuclei) of 1:500 dilution was prepared in 5% NGS and tissues were then incubated for 2 hrs at room temperature.

Immunohistochemistry on lymph gland was performed with the following primary antibodies: mouse anti-P1 (1:100, I. Ando), rabbit anti-Pxn (1:2000, J. Shim), rabbit anti-PPO (1:1000, H. Müller), rabbit anti-Dopamine (1:10,000, Abcam 6427), mouse anti-TH (1:100, Immunostar #22941), rabbit anti-Ddc (1:1000, Jay Hirsh), rat anti-DAT (1:1000, Millipore MAB369), rabbit anti-pH3 (1:100, CST #3642S), rabbit anti-Caspase3 (1:200, CST #9661S), rabbit anti-GFP (1:100, Abcam ab6556).

The following secondary antibodies were used at 1:500 dilutions: FITC, Cy3 and Cy5 (Jackson ImmunoResearch Laboratories) and Alexa Flour 546, 647 (Invitrogen) that was mouse, rabbit or rat specific. Nuclei were stained and visualized using Hoechst 33342 (Sigma B2261). Samples were mounted with Vectashield (Vector Laboratories). A minimum of ten independent biological replicates were analysed from which one representative image is shown.

### Dopamine staining of lymph gl and tissues

For detection of intracellular dopamine levels in the lymph gland, *Drosophila* larvae of different developmental timings (as per experimental requirement) were dissected in 1X PBS. This was followed by fixation in 4% formaldehyde for 20-25 mins and then stringent washes in 0.3% PBT (1X PBS + 0.3% Triton X 100) for 45 mins (3 washes for 15 mins each). The tissues were then incubated in 5% NGS for 45 mins. Dopamine antibody was then diluted in 5% NGS at a 1:10,000 dilution which was subsequently used to stain the lymph gland tissues overnight (15-18hrs) in 4°C. The secondary antibody treatment was conducted using the standard protocol described above.

### Imaging

Immuno-stained lymph glands images were acquired using Olympus FV3000 confocal microscopy under 40X oil-immersion objective with or without digital zooming (mentioned for the relevant experiment in the figure legends). It was made sure that the settings for acquisition were unaltered for different genotypes for each experimental set.

### Quantification of lymph gland phenotypes

All images were quantified using ImageJ software (NIH, USA) software (available at imagej.nih.gov/ij). The detailed methodology for analysis of various parameters is described as follows.

### Lymph gland and zone area analysis

The lymph gland and medullary/cortical zone area analysis was conducted as described in Shim et al., 2012. Specifically, middle two confocal Z-stacks were merged and the lymph gland area was estimated by outlining the primary lobe with the freehand selection tool in ImageJ followed by selecting the ‘analyse-measure’ option. This was also done for respective zones where Dome/Tep4 was used to estimate the progenitor area while P1 was used for the differentiated zone analysis. The zone areas is represented in percentage values, normalised to lymph gland area, in the graphs. In order to estimate the Dome/P1 overlap area, a single middle stack of the acquired confocal image was selected that best represented the progenitors and differentiated cells. The total lymph gland, Dome positive and P1 positive areas was measured by outlining (as mentioned above). The outlined region of interest (ROIs) of Dome and P1 areas marked were selected and then subjected to ‘More→ AND’ command in the ROI manager of ImageJ to give the overlap area. This was normalised to total lymph gland area and plotted as %Dome/P1 overlap. Controls were analysed in parallel to the tests every time. A minimum of 10 animals were analysed each time and the experiment was repeated at least three times.

### Crystal cell and pH3 positive nuclei analysis

The crystal cell population was assessed by counting the total number of PPO positive cells across the confocal Z stacks of the lymph gland and represented as crystal cells per lobe. Similarly, pH3 positive nuclei was counted across the Z stacks and normalised to total number of nuclei of the primary lobe and represented as ‘mitotic index’. A minimum of 10 animals were analysed each time and the experiment was repeated at least three times.

### Total nuclei count analysis

To count total nuclei, we adapted the spot detection methodology available in IMARIS, used for similar analysis in Sharma et al., 2019 and Rodrigues et al., 2021. Briefly, the confocal Z stacks of the lymph gland tissue were merged and the primary lobe was outlined using freehand selection tool. Following this, the remaining tissue like the ring gland, dorsal vessel, secondary lobes were cleared using the clear outside tool (under ‘Edit’ option). The nuclei of the primary lobe were then carefully thresholded using ‘Image-Adjust-Threshold’ option and applied to obtain an 8-bit image. The resulting image was then used for watershed (under ‘Process-Binary’ command) followed by particle analysis where the parameter used for nuclei size was (3-infinity) and circularity (0.04-1.00). The obtained ROIs as a result were then overlayed on the original image to avoid under/over counting of nuclei. A minimum of 10 animals were analysed each time and the experiment was repeated at least three times.

### FUCCI area analysis

To estimate the G1, S G2/M population in the progenitor pool, middle 2 Z stacks were merged, that best represented the medullary zone (Shim et al., 2012), of the acquired confocal image of the lymph gland at a specific development time point. After this, the channels of the merged image were separated to analyse the green and red channels, individually. For this, FUCCI reporter positive cells (green/red) were thresholded followed by ‘watershed’ and ‘analyse particles’ command to obtain the number of green and red positive cells, separately. To estimate the yellow population, the ROIs of the green positive cells were overlayed on the red channel image and the “green only” cell was counted manually. This “green only” value represents number of cells in G1 phase which was then eliminated from the total green positive cells counted, to obtain the yellow cells that represents G2/M population. The “red only” which represents S phase was similarly acquired by eliminating yellow cells from the total red cells calculated. These values were then normalised to the total FUCCI positive cells of the merged stacks and represented as %G1, %S or %G2/M population.

### Intensity analysis

The mean intensities in lymph gland tissues was calculated as described in Madhwal et al., 2020, Sharma et al., 2019. Briefly, the middle 2 stacks of the lymph gland images were selected, the area to be measured per lobe was defined using the freehand selection tool and the mean intensity measurement was then conducted using the measure tool. The relative fold change in intensities per lobe was calculated using mean intensity values. For all intensity quantifications, the laser setting for each individual experimental set-up was kept constant. Controls were analysed in parallel to the tests every time. A minimum of 10 animals were analysed each time and the experiment was repeated at least three times. The quantifications shown are for all the sets analysed.

### Statistical Analysis

All the statistical tests for the respective experiments were carried out using GraphPad Prism 9 and Microsoft Excel 2019 software. Either Student’s t-test or Mann-Whitney U test depending on the distribution of the data set acquired. The images represented have processed in ImageJ and Adobe Illustrator (2020).

## Acknowledgements

We thank I. Ando for the P1 antibody, Jay Hirsh for Ddc antibody, Utpal Banerjee and Lolitika Mandal for help with fly reagents, Bloomington Drosophila Stock Center (BDSC), Vienna Drosophila Resource Center (VDRC) and FlyBase for fly stocks. We acknowledge NCBS, CCAMP for CIFF and fly facility. We thank Apurva Sarin, Dasaradhi Palakodeti and inStem colleagues for helpful discussion and comments on the manuscript. Due to space limitations, we apologize to our colleagues whose work is not cited. This study was supported by the DBT-Center of Excellence grant BT/PR13446/COE/34/30/2015, DST-ECR ECR/2015/000390, 000390DBT-IYBA 2017, CEFIPRA and DBT Ramalingaswami Re-entry Fellowship to TM. AK is a Graduate Student at inStem, in the Mukherjee lab.

